# Microbial community shifts reflect losses of native soil carbon with pyrogenic and fresh organic matter additions and are greatest in low-carbon soils

**DOI:** 10.1101/2020.08.12.249094

**Authors:** Thea Whitman, Silene DeCiucies, Kelly Hanley, Akio Enders, Jamie Woolet, Johannes Lehmann

## Abstract

Soil organic carbon (SOC) plays an important role in regulating global climate change, carbon and nutrient cycling in soils, and soil moisture. Organic matter (OM) additions to soils can affect the rate at which SOC is mineralized by microbes, with potentially important effects on SOC stocks. Understanding how pyrogenic organic matter (PyOM) affects the cycling of native SOC (nSOC) and the soil microbes responsible for these effects is important for fire-affected ecosystems as well as for biochar-amended systems. We used an incubation trial with five different soils from National Ecological Observatory Network sites across the US and ^13^C-labelled 350°C corn stover PyOM and fresh corn stover OM to trace nSOC-derived CO_2_ emissions with and without PyOM and OM amendments. We used high-throughput sequencing of rRNA genes to characterize bacterial, archaeal, and fungal communities and their response to PyOM and OM. We found that the effects of amendments on nSOC-derived CO_2_ reflected the unamended soil C status, where amendments increased C mineralization the most in low-C soils. OM additions produced much greater effects on nSOC-CO_2_ emissions than PyOM additions. Furthermore, the magnitude of microbial community composition change mirrored the magnitude of increases in nSOC-CO_2_, indicating a specific subset of microbes were likely responsible for the observed changes in nSOC mineralization. However, PyOM responders differed across soils and did not necessarily reflect a common “charosphere”. Overall, this study suggests that soils that already have low SOC may be particularly vulnerable to short-term increases in SOC loss with OM or PyOM additions.

**Importance:** Soil organic matter (SOM) has an important role in global climate change, carbon and nutrient cycling in soils, and soil moisture dynamics. Understanding the processes that affect SOM stocks is important for managing these functions. Recently, understanding how fire-affected, or “pyrogenic” organic matter (PyOM) affects existing SOM stocks has become increasingly important, both due to changing fire regimes, and to interest in “biochar” – pyrogenic organic matter that is produced intentionally for carbon management or as an agricultural soil amendment. We found that soils with less SOM were more prone to increased losses with PyOM (and fresh organic matter) additions, and that soil microbial communities changed more in soils that also had greater SOM losses with PyOM additions. This suggests that soils that already have low SOM content may be particularly vulnerable to short-term increases in SOM loss, and that a subset of the soil microbial community is likely responsible for these effects.

## Introduction

Soil organic matter (SOM) supports a wealth of benefits in soil systems, including providing organic nutrients, binding toxic compounds, increasing soil water holding capacity, and storing soil organic carbon (SOC). Globally, soils hold large stocks of carbon (C) – twice the amount of C held in living biomass or in the atmosphere (1). Understanding the processes that control the stocks and fluxes of C in and out of the soils is thus essential for mitigating climate change, as well as for sustainable agricultural management (2). Recently, the importance of understanding the role of pyrogenic, or fire-affected, organic matter (PyOM) (*sensu* Zimmerman and Mitra (3)) in contributing to SOC stocks has become increasingly salient. PyOM plays an important role in contributing to soil carbon stocks, particularly in fire-affected ecosystems (4), and can represent over 60% of total SOC (5). Its persistence in soils has led to interest in its role in offsetting the climate impacts of natural wildfires (4) as well as the possibility of its intentional production for the stabilization of organic matter (OM), in which case it is often referred to as “biochar” (6, 7). However, in order to quantify its net effect on C stocks and fluxes, it is essential to understand not only the persistence of pyrogenic C (PyC) itself, but also its effect on the native SOC (nSOC) present before PyC additions.

After interest was sparked in the potential of PyC for climate change mitigation just over a decade ago, alarm bells were sounded about the possibility of its addition to soils resulting in increased loss of nSOC and increased CO_2_ emissions (8–10). These observations sparked a flurry of research into the potential interactions between added PyC and nSOC. This research was important, because if PyOM additions are to be used for climate change mitigation, it must not be offset by increased nSOC losses. Initial investigations revealed a range of responses, spanning from large increases in nSOC mineralization to large decreases in SOC mineralization with PyOM additions (11–13). (Although the term “priming” (14) is widely-used to describe this phenomenon, due to broad interpretations of the term (15), we will refer to “increased or decreased mineralization”; even though less concise, this method- and process-agnostic term will help ensure clarity and avoid prior expectations of what the term “priming” implies.) Research over the past decade has progressed beyond observation of the phenomenon to systematic investigations of the mechanisms underlying these interactions (16–18), while the conclusions from meta-analyses have strengthened as the total number of studies of PyOM-SOC interactions has steadily increased (19–22).

The above-cited meta-analyses provide a robust overview of recent advances in the literature. Briefly, current understanding of mechanisms underlying interactive effects of PyOM additions on SOC mineralization includes the following observations (19–22): (1) In general, when changes in mineralization do occur, net increases in nSOC mineralization tend to be limited to the earlier stages of incubations or field studies, while net decreases in nSOC mineralization often emerge later. (2) It is essential to consider the specific properties of PyOM and the soil to which it is applied together. Properties such as pH, total nSOC content, nutrient status, and texture or particle size are important determining factors of the net C effects of PyOM additions on nSOC. (3) Specific researcher-determined conditions of the study can significantly determine the effects of interest. This is particularly true for moisture and duration of the experiment. Although the above factors make it challenging to collectively develop a predictive understanding of interactions between SOC and PyOM mineralization, it is important to design experiments explicitly to test for and quantify the relative importance of specific mechanisms. In this spirit, in this study, we sought to investigate short-term increases in SOC mineralization with PyOM amendments. Although numerous studies have now observed net decreases in SOC mineralization with PyOM amendments over the long term, characterization of the mechanisms that underpin both of these phenomena will help us develop appropriate models for predicting long-term effects into the future (23, 24). For example, in a C cycling model designed to predict the long-term effects of PyOM on C stocks (23), the assumption is that the dominant mechanism of decreased SOC mineralization is sorption of SOC by the PyOM, which is represented in the model by decreasing the fraction of SOC that is partitioned to the more rapidly-cycling pool. However, the assumption for increased SOC mineralization is that the dominant mechanism is increased microbial activity, which is represented in the model by increasing the rate at which nSOC is mineralized. These assumptions create a model structure that helps drive the model’s predictions of long-term net decreases in nSOC mineralization with PyOM additions. Although increases in nSOC mineralization rates after PyOM additions seem to be limited to short- and medium (<2 year) timelines (21, 22), we wanted to investigate these short-term effects, since they pose the greatest risk of unintended consequences for nSOC stocks during intentional PyOM additions as biochar for C management or for increased nSOC losses due to PyOM inputs after wildfires.

Commonly proposed mechanisms for short-term increases in nSOC mineralization with PyOM additions can be broadly grouped in two: (A) *co-metabolism*: easily mineralizable PyOM fractions increase microbial activity, resulting in additional decomposition of SOC; (B) *stimulation*: PyOM additions may result in changes to the soil chemical or physical environment that generally favour increased microbial activity, such as more optimal pH, nutrient, oxygen, or water conditions (19, 20). In addition, *community composition shifts* could also help explain these phenomena (25). It is possible that PyOM additions could induce changes to the microbial community composition that shift the community toward taxa that favour different sources of organic matter, or process organic matter differently – *e.g.*, organisms with different carbon use efficiencies (CUE) (18). Finally, researchers often distinguish these effects from “apparent priming” – when total CO_2_ emissions from soil increase, but this increase is not accompanied by increases in nSOC losses (15). Rather, the increase is attributed to increased turnover of soil microbial biomass. While the effects included under *stimulation* are essential to understand in order to predict SOC fluxes, they are – mechanistically – comparably straightforward: researchers have long studied the effects of changing moisture or oxygen on SOC fluxes. If we are able to quantify the degree to which PyOM additions to soil change these properties, we will be on our way to predicting their effects on nSOC cycling. However, the effects included under *co-metabolism* and *community composition shifts* are generally less well-characterized, and it is these mechanisms that we specifically sought to investigate in this study.

While research into the mechanisms behind changes in SOC mineralization with PyOM additions has grown substantially over the last decade, our understanding of which microbes respond to PyOM additions, and the reasons for their response, has somewhat lagged behind, particularly for fungi. As an exception to this, the recent investigation by Yu *et al*. into the effects of PyOM on SOC mineralization included an assessment of bacteria and fungi, using high-throughput sequencing, through which they identified that the relative abundance of fungal classes *Sordariomycetes* and *Tremellomycetes* were significantly positively correlated with increases in SOC mineralization after 40 days of incubation (26). In our recent review of PyOM effects on soil bacterial communities (27), we re-analyzed papers published before 2018 that had publicly accessible data and used Illumina high-throughput sequencing of the 16S ribosomal RNA gene to characterize soil bacterial communities (25, 28–32). Using the same approach to reanalyze all datasets, we found the following: (A) although most communities were significantly altered by the addition of PyOM, rather than creating a “charosphere”-dominated community (33, 34), PyOM-amended soil bacterial communities resembled their corresponding unamended soil communities more closely than they resembled different soils that had also been amended with PyOM; (B) phylum-level responses to PyOM additions were not consistent across different soil and PyOM combinations – *i.e.*, taxonomic level is generally too broad to make meaningful conclusions about soil bacterial responses to PyOM; and (C) a small number of taxa were identified as being PyOM-responders in more than one study, most of which came from the phyla *Actinobacteria* and *Proteobacteria* (27). Based on these findings, we would suggest that the field is still too nascent to make broad generalizations about any kind of consistent effect of PyOM on microbial communities, and hope that continuing to blend functional measurements with microbial response data will help to identify which specific microbes might be responsible for changes in nSOC mineralization with PyOM additions, while also generally increasing our understanding of which microbes respond to PyOM additions and why.

In this study, we had two research questions, with alternate hypotheses for each. Our first question was, are soils with less SOC or less mineralization more prone to stimulation by PyOM additions? Our primary hypothesis was that soils with less nSOC mineralization are more likely to experience increased mineralization with PyOM additions via *co-metabolism*. Our rationale was that these microbial communities are more likely to be C-limited, and the addition of (the easily-mineralizable fraction of) PyOM could alleviate this constraint (16). On the other hand, our alternate hypothesis rationalized the opposite: soils with less nSOC or less mineralization may be less likely to experience increased short-term mineralization with PyOM additions. This could occur if the microbial communities were limited by mineral nutrients. If PyOM additions alleviated this constraint via *stimulation*, microbial communities in soils with more mineralizable OC might be better able to take advantage of this subsidy. Our second question was, do soil microbial communities reflect changes in nSOC mineralization with PyOM additions? Our primary hypothesis was that there would be larger changes to the microbial community in the soils where PyOM additions increased nSOC mineralization, while microbial communities in soils that did not experience increased nSOC mineralization would not change as much. The rationale was that groups of microbes that respond positively to PyOM additions may be the same groups that are responsible for increased nSOC mineralization with PyOM additions, so a stronger shift toward these groups may accompany a stronger effect on nSOC mineralization. Our first alternate hypothesis was that PyOM might change microbial communities similarly in all soils – if PyOM additions had a very strong effect on the microbial community composition, creating a consistent “charosphere” community, differences from one soil to the next might be too subtle in comparison to detect. Our second alternate hypothesis was that we might not see substantial community shifts at all with PyOM additions. Although previous studies have seen significant changes to microbial communities with PyOM additions (27), these studies have often added extremely high amounts of PyOM. When applied at environmentally relevant rates, while PyOM additions may provide an additional C source, it may be relatively small in comparison to available nSOC, and any effects of PyOM additions on soil water-holding capacity, pH, or nutrient availability may not be large enough to clearly affect the soil microbial community composition.

In order to investigate these questions, we selected five different soils with a range of SOC stocks and associated mineralization rates and applied ^13^C-labelled PyOM produced at 350°C from corn (*Zea mays* L.) stover as well as the original ^13^C-labelled corn stover OM, tracing CO_2_ fluxes continuously over one month, and characterizing the response of the soil microbial community using high-throughput sequencing of the bacterial and archaeal (16S) and fungal (ITS2) communities.

## Materials and Methods

### Experimental overview

We incubated five contrasting soils from sites across the United States (Table 1), adding ^13^C-labelled corn stover (“OM”), PyOM produced at 350 °C from the same corn stover (“PyOM”), or no additions (“Soil”). We monitored CO_2_ fluxes over four weeks and used stable isotope partitioning to separate CO_2_ emissions into SOC- and amendment-derived pools. We characterized the microbial (bacterial/archaeal and fungal) communities after 24 hours, 10 days, and 26 days using ribosomal RNA gene sequencing.

**Table 1.** Studied soils and their properties

| Source | Soil Type | C (%) | N (%) | Ca (mg kg<sup>-1</sup>) | Mg (mg kg<sup>-1</sup>) | Na (mg kg<sup>-1</sup>) | K (mg kg<sup>-1</sup>) | pH |
| --- | --- | --- | --- | --- | --- | --- | --- | --- |
| Laupahoehoe, HI | Typic Hydrudand | 33.1 | 2.0 | 492 | 137 | 29 | 86 | 5.0 |
| Caribou Creek-Poker Flats, AK | Pergelic Cryaquept | 9.4 | 0.4 | 540 | 68 | 16 | 20 | 5.0 |
| Ithaca, NY | Typic Fragiudept | 4.6 | 0.4 | 1219 | 202 | 87 | 119 | 5.1 |
| Onaqui, UT | Xeric Haplocalcid | 2.6 | 0.1 | 5619 | 245 | 28 | 511 | 8.3 |
| Ordway-Swisher Biol. Station, FL | Lamellic Quartzipsamment | 0.5 | 0.02 | 77 | 14 | 11 | 8 | 5.2 |

### Soil descriptions

Soil properties are described in Table 1. Four of the soils were collected from National Ecological Observatory Network (NEON) sites following NEON protocols in 2009-2010, as part of a NEON prototype study, from the top 0-0.1 m of the A horizon (35). We added a local non-NEON site in which we had previously investigated PyOM effects on SOC cycling and microbial communities (the Fragiudept / NY site). Each sample was from a single core, except for the Cryaquept and Fragiudept, which were from composited cores. Samples were stored at - 80 °C until experimental initiation, except for being shipped overnight to Ithaca, NY on dry ice. Clearly, this treatment would be expected to have an effect on the specific soil community composition. Due to logistical constraints in collecting fresh soils from all sites, we worked with frozen samples. Thus, we would expect our findings for these soils with respect to the dominant mechanisms at work to remain applicable in other systems, while the specific effects (*e.g.*, baseline abundances of individual taxa or absolute magnitude of CO_2_ fluxes) should not be directly translated to natural ecosystems.

### Corn stover and PyOM amendment production

^13^C pulse-labelled corn (*Zea mays* (L.)) shoot biomass was grown, ground (<2 mm), and pyrolyzed at 350 °C under Ar gas in a modified muffle furnace as previously described in detail (36). Amendment properties are reported in Supplemental Table 1.

### Incubation setup and monitoring

Frozen samples were thawed, sieved <2 mm, and air-dried at room temperature, until mass stabilized with losses changing by less than 1% per day. A sub-sample was rapidly dried at 70 °C in a drying oven and used to determine moisture-holding capacity individually for each soil, with each amendment, in order to ensure that all samples are at equivalent moisture levels, given that amendments might affect water holding capacity. To do this, we weighed the soil samples (amended or unamended) into a PVC tube with a screen covered by a moist filter paper at the bottom. The tubes were placed in a container and water was slowly added to the container until the samples were saturated and the level of the water was level to the surface of the soil. The saturated soils were let stand overnight. In the morning, they were removed from the water bath, and allowed to drain freely overnight, covered in parafilm. The mass of water remaining in the soil was taken to represent “field capacity” (FC), with a target moisture value for incubation of 65% FC. We also calculated the final moisture content of the air-dried soil, to enable us to calculate the water required to reach this value for the incubations.

We prepared separate incubation vials for each treatment to be sampled at each timepoint. This was done so that we could destructively sample them completely, in order to ensure representative sampling, and so that we could be certain of the masses in the remaining incubation jars. For vials with amendments, we added OM at 3% by mass, and added PyOM on a pre-pyrolysis mass basis, which resulted in a 0.99% by mass addition. *I.e.*, we added the mass of PyOM that would have remained if we used the same amount of initial biomass to produce PyOM, essentially asking the systems-level question, “What might the fate of this biomass be?”. Based on our expectations for CO_2_ flux rates from previous experiments, we determined that we would require 1 g soil per incubation for the high-organic matter soils (Typic Hydrudand and Perigelic Cryaquept), and 5 g per incubation for the lower organic matter soils, in order to make sure that CO_2_ fluxes remained within the optimal range for our instrumentation setup. For the 24 h timepoints, we used 2 g of soil. Each jar – amended and unamended – was stirred to mix. The experiment was initiated (t_0_) for each jar when water was added to bring it up to 65% FC. At wet-up, each jar received water drop-wise, to gradually bring it up to the target moisture level. The vial for the 24-h timepoint was incubated at 30 °C for 24 h in Mason jars with 20 mL DIW in the bottom to maintain a moist environment, and was then destructively sampled for microbial community composition after exactly 24 h, by collecting the entire sample in a Whirl-Pak bag. The sample was immediately frozen at −80 °C and stored until DNA extraction, except for overnight shipment on dry ice to Madison, WI. The two vials for the two later timepoints – 10 d and 26 d – were placed in the same quart-size Mason jar, along with 20 mL DIW in the bottom of the Mason jar to maintain a moist environment. The Mason jar was then sealed with a lid with tubing connected to the gas monitoring system. Because each full measurement cycle on the gas monitoring system takes 20 minutes, one experimental treatment was wet up every 20 minutes, taking care to attach it to the gas monitoring system at the corresponding time. The jars were automatically sampled using a custom-built multiplexer system (See (17) for details), connected to a Cavity Ring-Down Spectrometer (Picarro G2201-I, Santa Clara, CA, USA) that measures CO_2_ concentrations and ^13^CO_2_/^12^CO_2_ isotopes. Measurements were made on a continuous monitoring cycle, which resulted in each Mason jar being measured about once a day. After 10 days, jars were opened, and one vial was randomly removed to be destructively sampled for microbial community characterization. Mason jars were removed and returned on a time cycle to ensure that each vial was sampled at the equivalent time since wet-up. After 26 days, the second vial was removed and destructively sampled for microbial community characterization.

### DNA extraction and sequencing

DNA extractions were performed for each sample and for the original materials (OM and PyOM), with one blank extraction for every 24 samples (identical methods but using empty tubes, all of which were sequenced). We used a DNEasy PowerLyzer PowerSoil DNA extraction kit (QIAGEN, Germantown, MD) following manufacturer’s instructions and bead-beating samples for 45 s at 6 m s^−1^ on a FastPrep 5G homogenizer (MP Biomedicals, Santa Ana, CA). Extracted DNA was amplified in triplicate PCR, targeting the 16S rRNA gene v4 region (henceforth, “16S”) with 515f and 806r primers (37), and targeting the ITS2 gene region with 5.8S-Fun and ITS4-Fun primers (38) with barcodes and Illumina sequencing adapters added as per (39) (all primers in Supplemental Tables 2-4). The PCR amplicon triplicates were pooled, purified and normalized using a SequalPrep Normalization Plate (96) Kit (ThermoFisher Scientific, Waltham, MA). Samples, including blanks, were pooled and library cleanup was performed using a Wizard SV Gel and PCR Clean-Up System A9282 (Promega, Madison, WI). The pooled library was submitted to the UW Madison Biotechnology Center (UW-Madison, WI) for 2×250 paired end (PE) Illumina MiSeq sequencing for the 16S amplicons and 2×300 PE for the ITS2 amplicons.

### Microbial community bioinformatics

For 16S reads (32k min, 208k max, 58k median total sequenced reads), we quality-filtered and trimmed, dereplicated and learned errors, assigned operational taxonomic units (OTUs), and removed chimeras, using dada2 (40) as implemented in R (mean 53% of initial reads remaining after full pipeline; 15k min, 180k max, 29k median total final reads). Taxonomy was assigned to the 16S reads using a naïve Bayes classifier (41) trained on the 515f-806r region of the 99% ID OTUs from the Silva nr 132 database(42) (Yilmaz *et al*., 2014) as implemented in QIIME2 (43). We removed any OTUs classified as chloroplasts or mitochondria. For ITS2 reads (21k min, 289k max, 62k median total sequenced reads), we first merged reads using PEAR (44), and then performed the same steps as described for 16S above (mean 50% of initial reads remaining after full pipeline; 6k min, 199k max, 32k median total final reads). Taxonomy was assigned to the ITS2 reads using the UNITE general release dynamic threshold database (02.02.2019) (UNITE, 2019) using a naïve Bayes classifier (41) as implemented in dada2 (40). We removed any OTUs that did not receive a classification at the phylum level in order to exclude any non-fungal ITS2 sequences. High-memory-intensive sequence processing steps were performed on the UW-Madison Centre for High Throughput Computing cluster (Madison, WI).

### Stable isotope CO_2_ flux partitioning

Respiration data were analyzed as per (17) using R version 3.6.1 (45). Sample respiration was partitioned between the amendment-derived CO_2_-C and soil-derived CO_2_-C using the following equations (46):

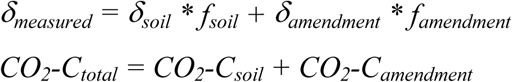

where δ represents the δ^13^C signature (with respect to the PeeDee Belemnite standard) of the total respired CO_2_-C (δ_measured_), the soil-derived CO_2_ (δ_soil_), or the amendment-derived CO_2_-C (δ_amendment_), and *f* represents the fraction of the total CO_2_-C derived from the soil (*f*_*soil*_) or the amendment (*f*_*amendment*_). δ^13^C of bulk PyOM (δ_PyOM_) or bulk OM (δ_OM_) was used as the amendment endmember for isotope partitioning. Soil isotope endmembers (δ_soil_) to be used in isotope partitioning were obtained daily using the average δ^13^C for CO_2_-C from control (unamended) treatments (See supplemental R scripts). We interpret values that do not overlap within a 95% confidence interval as being significantly different.

### Microbial community analyses

We worked primarily in Jupyter notebooks, with phyloseq (47), ggplot (48), and dplyr (49) being instrumental in working with the data in R (45). We compared community composition across samples using Bray-Curtis (50) dissimilarities on Hellinger-transformed relative abundances (51), which we represented using NMDS ordinations. We tested for significant effects of soil site, days of incubation, amendment, and interactions between soil and day, and soil and amendment using a permutational multivariate ANOVA (PERMANOVA; the adonis function in vegan (52). We identified OTUs that were differentially abundant (significantly enriched in amended soils as compared to control soils) within each soil type and amendment, testing only taxa that represented at least 0.01% of the mean total community for that soil using the R package corncob (53). We analyzed the two later timepoints together, while controlling for timepoint and controlling for differential variance, using a Wald test and correcting p values to yield a false discovery rate of less than 0.05 within each soil type and amendment.

### Data availability

Sequencing data are available in the NCBI SRA under accession numbers XXX. Code used to analyze data and generate figures in this paper is available at github.com/TheaWhitman/NEON_PyOM.

## Results

nSOC-derived CO_2_ emissions were greatest in the soils with the most total SOC (the Hydrudand and Cryaquept) and lowest in the soils with less total SOC (the Haplocalcid, Fragiudept, and Quartzipsamment) (Figure 1). PyOM additions increased cumulative nSOC-derived CO_2_ emissions by 55% for the Quartzipsamment soil (FL) only, while OM additions increased cumulative nSOC-derived CO_2_ emissions by 44% for the Haplocalcid (UT), by 126% for the Fragiudept (NY), and by 170% for the Quartzipsamment (FL) soils (Figures 1 and 2). These effects were generally largest earlier in the incubation periods, although the significant effects persisted throughout the full 26 days for the Quartzipsamment.

**Figure 1.**
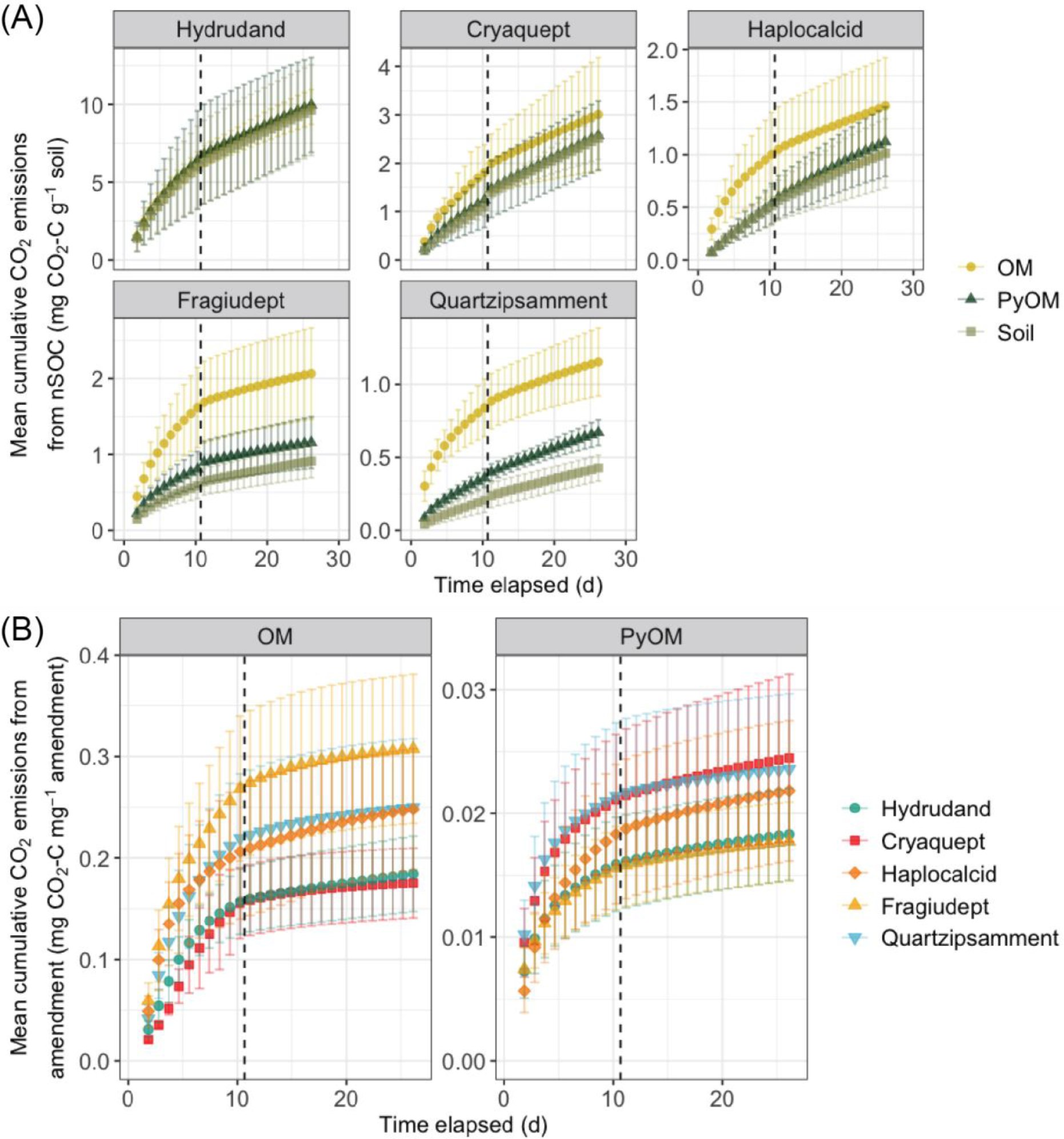
(a) Mean cumulative nSOC-derived CO_2_ emissions over time for each soil, with organic matter (OM; yellow circles) additions, pyrogenic organic matter (PyOM; dark green triangles) additions, or no additions (Soil; pale green squares). Error bars represent ±1.96SE (95% confidence interval). Dashed line indicates sampling point for mid-incubation harvests. N=4. Note different scales on the y-axes. (b) Mean cumulative amendment-derived CO_2_ emissions over time. Error bars represent ±1.96SE (95% confidence interval). Dashed line indicates sampling point for mid-incubation harvests. N=4. Note different scales on the y-axes.

**Figure 2.**
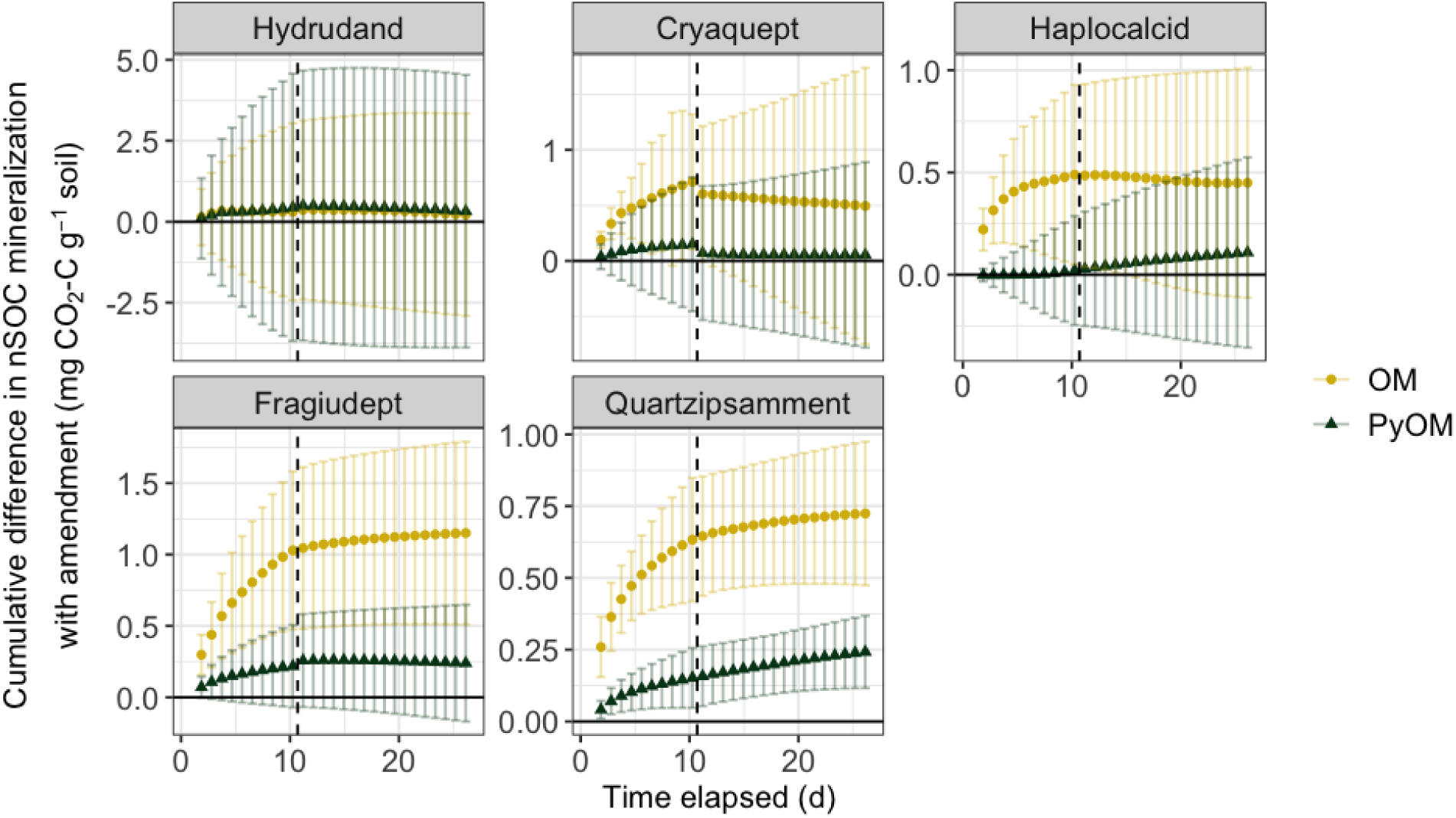
Mean cumulative difference in nSOC-derived CO_2_ emissions in amended soils as compared to unamended soil over time for each soil, with organic matter (OM, red circles) additions and pyrogenic organic matter (PyOM, orange triangles) additions. Error bars represent ±1.96SE (95% confidence interval). Dashed line indicates sampling point for mid-incubation harvests. N=4. Note different scales on the y-axis.

For the full dataset, bacterial community composition was significantly affected by soil site, days of incubation, amendment, and interactions between soil site and day, and soil site and amendment (PERMANOVA, p<0.001 for all effects; Supplementary Table S5; Supplemental Figure S1). When the soils were analyzed individually (Figure 3A), days of incubation and amendment were all significant predictors of bacterial community composition (PERMANOVA, p<0.02), except for the Hydrudand, where only days of incubation were significant (Supplementary Table S6). The effects of amendments were least pronounced in the Hydrudand from Hawaii and the Cryaquept from Alaska, and most pronounced for the Quartzipsamment from Florida (Supplementary Table S6).

**Figure 3.**
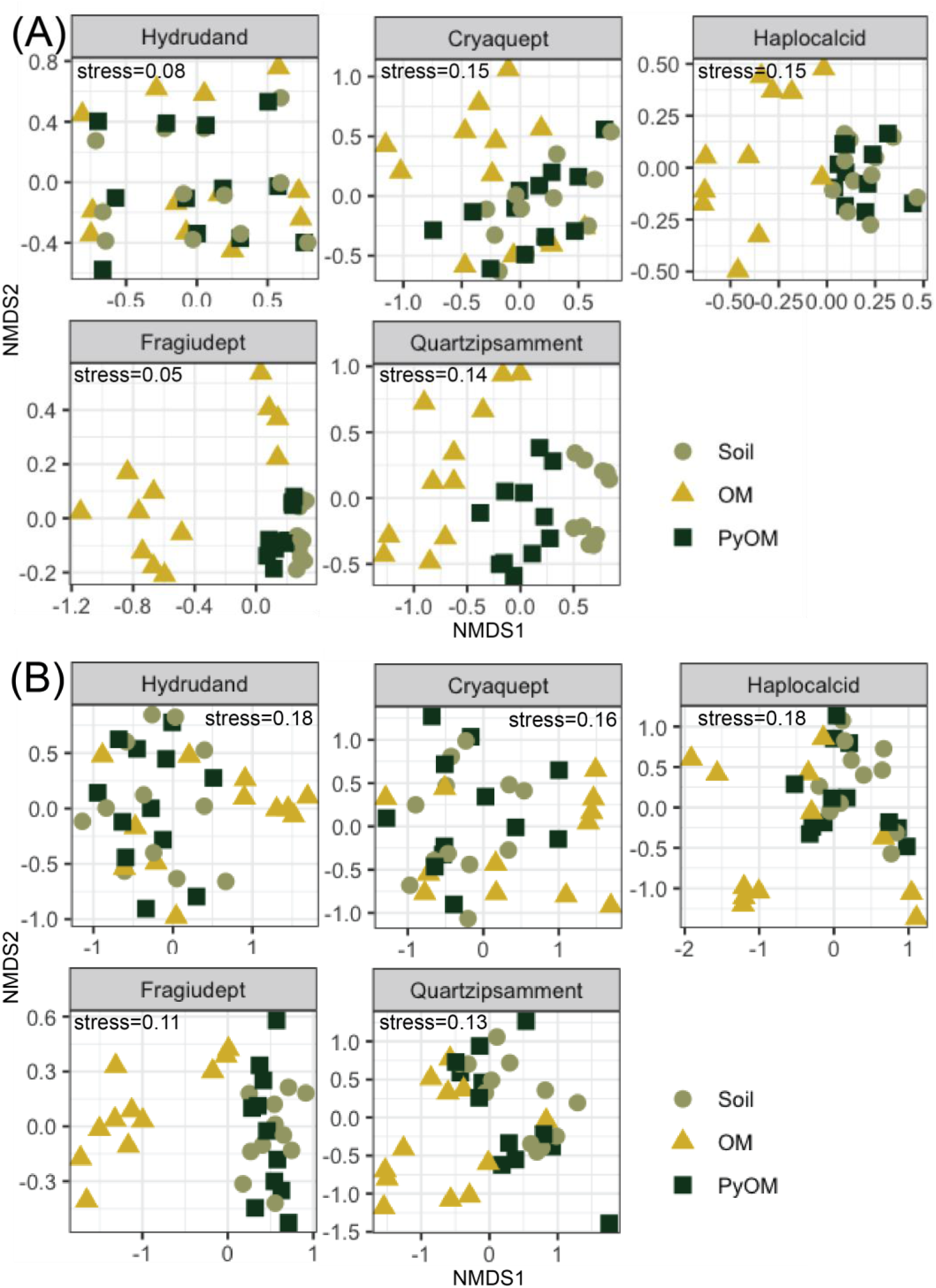
Non-metric multidimensional scaling plot of Bray-Curtis distances between soil microbial communities (Hellinger-transformed relative abundances) at all three timepoints (not distinguished on figure) for each soil. Shapes indicate whether organic matter (OM, yellow triangles), pyrogenic organic matter (PyOM, dark green squares), or nothing was added (Soil, light green circles). (A) Bacteria and Archaea (16S) k=2, stress_Hydrudand_=0.08, stress_Cryaquept_=0.15, stress_Haplocalcid_=0.15, stress_Fragiudept_=0.05, stress_Quartzipsamment_=0.14. N=4 for each timepoint, except Haplocalcid on day 10 and Quartzipsamment on day 26, where N=3; ordinations were performed individually for each soil type. (B) Fungi (ITS2) k=2, stress_Hydrudand_=0.18, stress_Cryaquept_=0.16, stress_Haplocalcid_=0.18, stress_Fragiudept_=0.11, stress_Quartzipsamment_=0.13. N=4 for each timepoint, except Fragiudept on day 1, where N=3; ordinations were performed individually for each soil type.

For the full dataset, fungal community composition was significantly affected by soil type/site, days of incubation, amendment, and interactions between soil and day, and soil and amendment (PERMANOVA, p<0.001 for all effects; Supplementary Table S7; Supplemental Figure S2). When the soils were analyzed individually (Figure 3B), amendment was a significant predictor of fungal community composition for all soils except the Cryaquept (PERMANOVA, p<0.007), and days of incubation were significant for the Hydrudand, Cryaquept, and Fragiudept (PERMANOVA, p<0.03) (Supplementary Table S8). The effects of amendments were most pronounced in the Fragiudept from New York (Supplementary Table S8).

The greatest changes in soil community composition (highest Bray-Curtis dissimilarity from unamended soil) upon amendment with PyOM or OM were associated with the greatest increases in nSOC-derived CO_2_ emissions (Figure 4).

**Figure 4.**
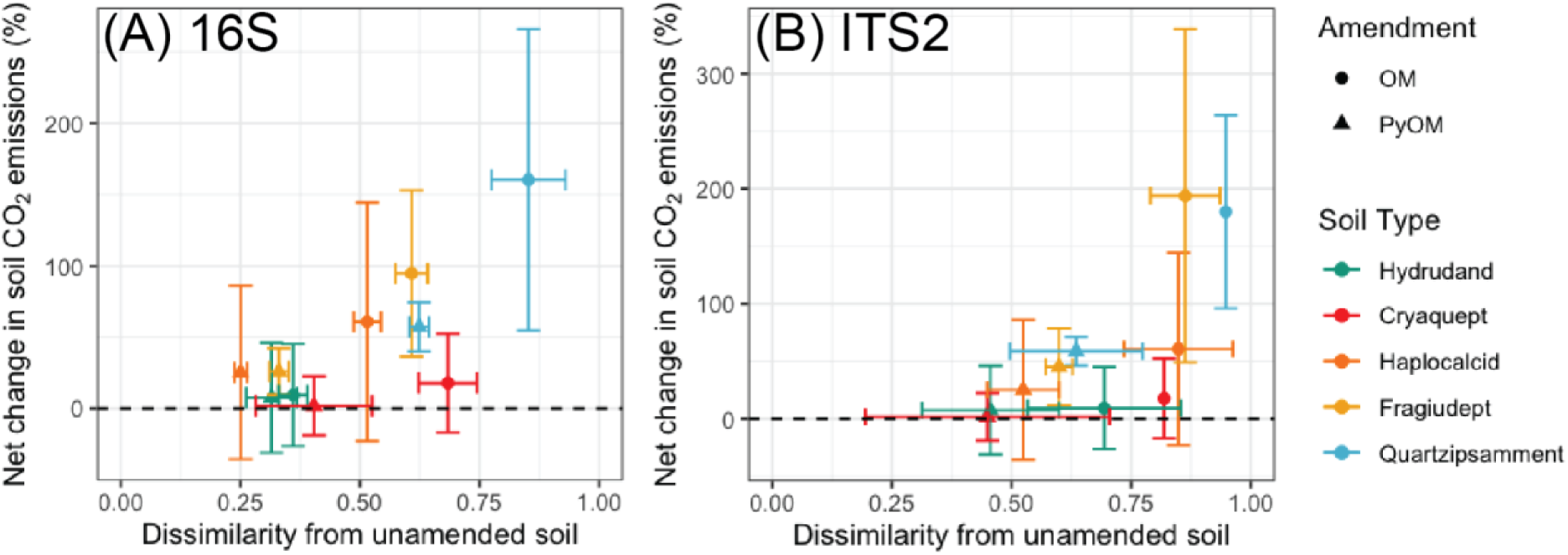
Net change in soil-derived CO_2_ emissions (%) vs. Bray-Curtis dissimilarity on Hellinger-transformed abundances from unamended soil for different soils and amendments at final timepoint (day 26). Circles represent samples amended with OM and triangles represent samples amended with PyOM. Error bars represent ± 1.96SE (95% confidence interval). (A) Bacteria and Archaea (16S); N=4, except Quartzipsamment, where N=3. (B) Fungi (ITS2); N=4.

Across all soils, we identified 258 16S OTUs that responded positively to OM amendments, and 162 OTUs that responded positively to PyOM amendments (Figure 5; Supplemental Table S9). Of these OTUs, 77 were responders to PyOM in at least one soil and to OM in at least one soil, or “common positive responders”. Genera with common positive responders in multiple soils included *Chthoniobacter* (9 OM-responsive OTUs across 3 soils and 5 PyOM-responsive OTUs across 2 soils), *Flavisolibacter* (3 OM-responsive OTUs in 1 soil and 3 PyOM-responsive OTUs across 2 soils), *Bacillus* (29 OM-responsive OTUs across all 5 soils and 6 PyOM-responsive OTUs across 2 soils), *Ammoniphilus* (3 OM-responsive OTUs across 2 soils and 2 PyOM-responsive OTUs across 2 soils), *Gemmatimonas* (3 OM-responsive OTUs across 2 soils and 6 PyOM-responsive OTUs across 3 soils), *Gemmata* (2 OM-responsive OTUs across 2 soils and 2 PyOM-responsive OTUs across 2 soils), *Anaeromyxobacter* (2 OM-responsive OTUs across 2 soils and 3 PyOM-responsive OTUs across 3 soils), *Microvirga* (18 OM-responsive OTUs across 3 soils and 8 PyOM-responsive OTUs across 3 soils), *Achromobacter* (1 OM-responsive OTUs and 2 PyOM-responsive OTUs across 2 soils), *Noviherbaspirillum* (8 OM-responsive OTUs across 3 soils and 5 PyOM-responsive OTUs across 2 soils), *Allorhizobium-Neorhizobium-Pararhizobium-Rhizobium* (1 OM-responsive OTU and 2 PyOM-responsive OTUs across 2 soils), and *Haliangium* (1 OM-responsive OTU and 2 PyOM-responsive OTUs across 2 soils). Only one fungal OTU – a *Spizellomyces* from the *Chytridiomycota* phylum – was identified as being a significant positive responder to PyOM (estimated log_2_-fold change of 6.3 in the Fragiudept), and no fungi were positive responders to OM over the timeframe of this study (Supplemental Table S10).

**Figure 5.**
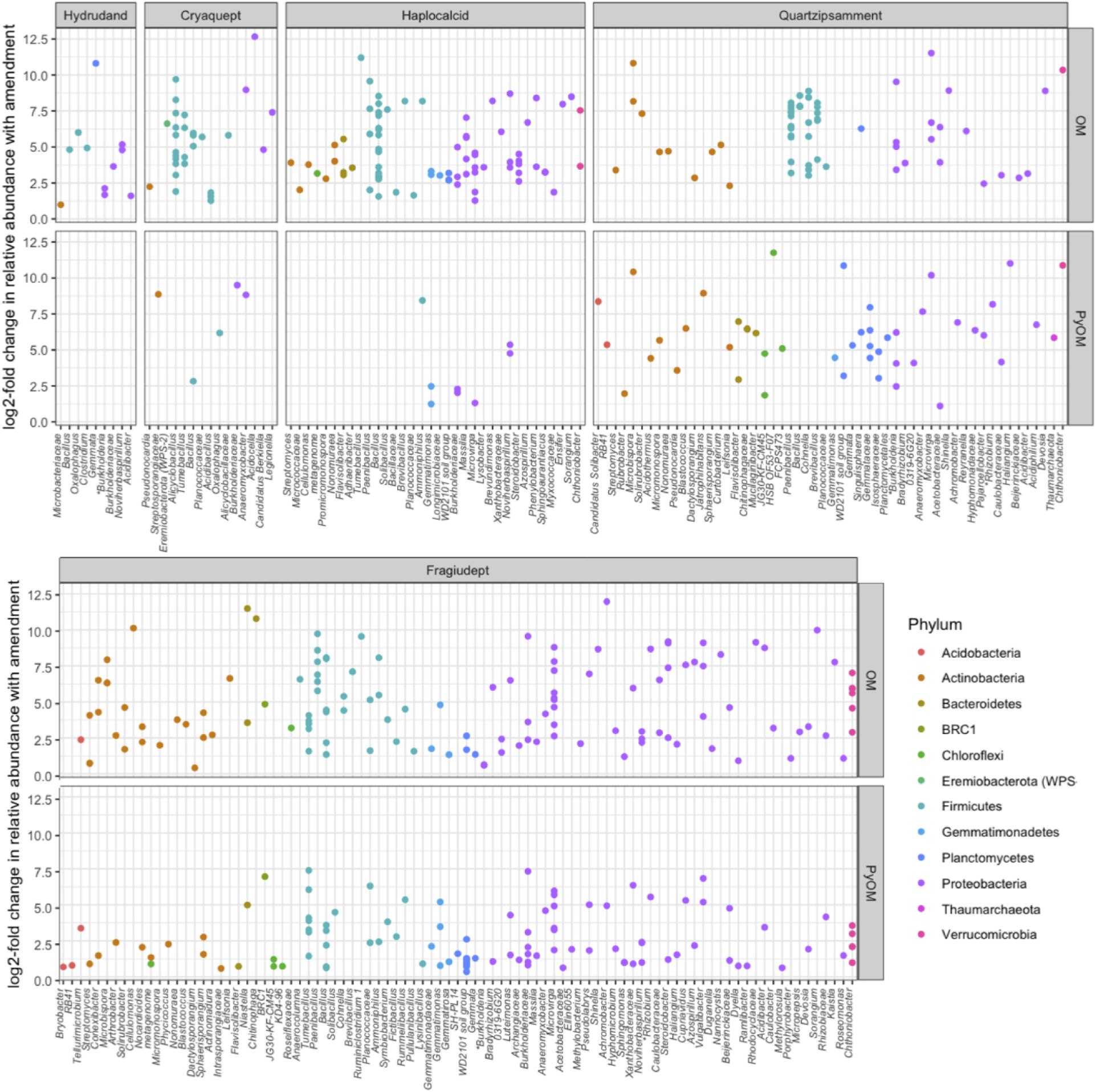
Differential abundance of bacterial and archaeal OTUs that are positive responders to OM or PyOM additions, as estimated using the “corncob” algorithm (53) and grouped by soil and finest taxonomic resolution available. Each point represents a single OTU. *Rhizobium label represents “Allorhizobium-Neorhizobium-Pararhizobium-Rhizobium” and *Burkholderia label represents “Burkholderia-Caballeronia-Paraburkholderia”.

With a few exceptions, bacterial taxa that responded positively or negatively to PyOM tended to also respond similarly to OM (Figure 6).

**Figure 6.**
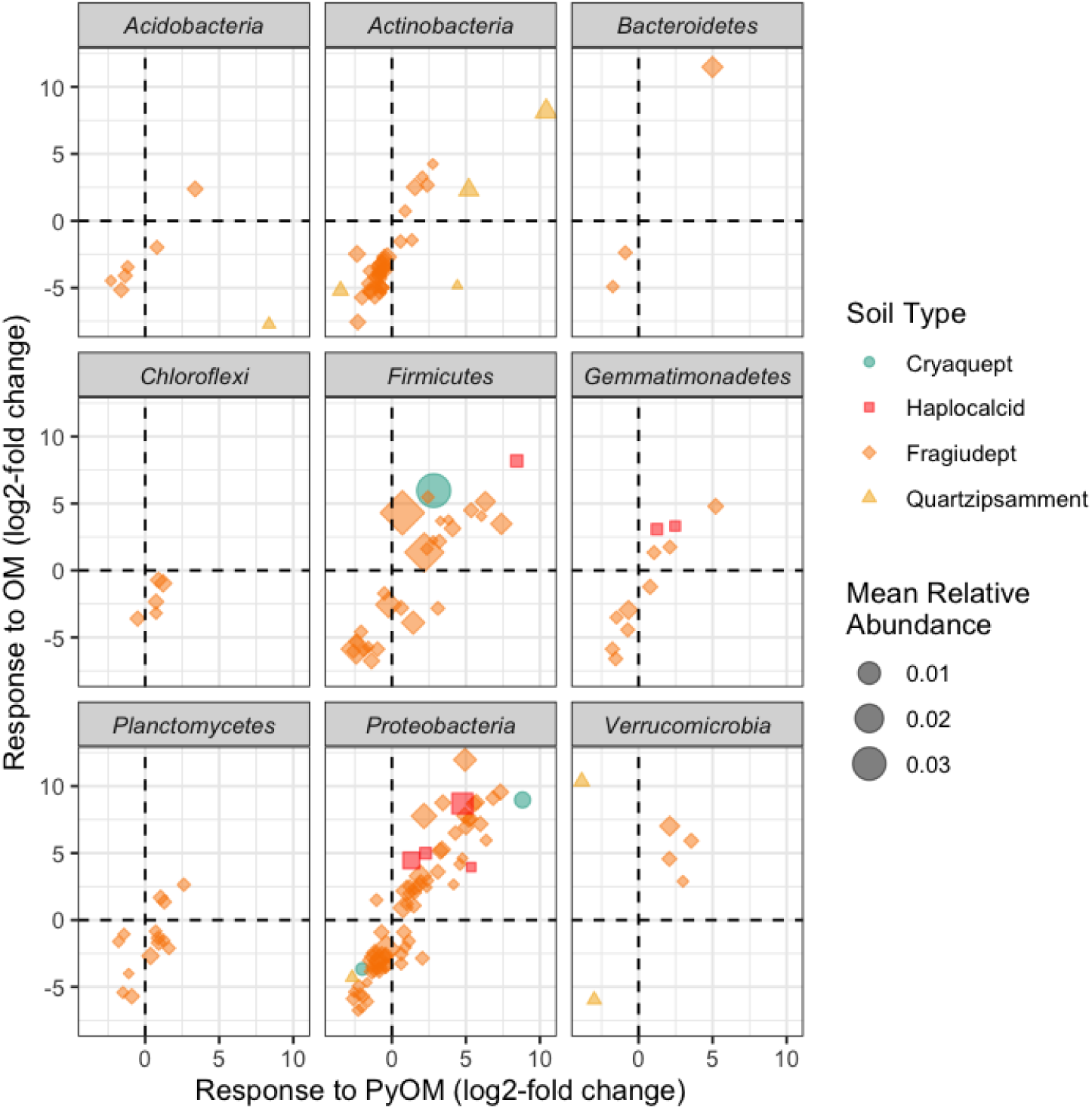
Response to PyOM vs. response to OM for bacterial OTUs that were present at a mean of at least 0.01% and for which there were sufficient observations to perform statistical testing in both OM- and PyOM-amended samples, as estimated using the “corncob” algorithm (53). Each point represents a single OTU from one soil, with color and shape indicating soil source, and size scaled by mean relative abundance within a soil, across all treatments, on days 10 and 26. Dashed lines indicate 0, or no change in relative abundance as compared to unamended soil.

## Discussion

### Effects of organic amendments on nSOC-derived CO_2_ reflect baseline soil C status

In response to our first question, our findings were consistent with our primary hypothesis: soils with lower baseline CO_2_ emissions experienced greater increases in nSOC mineralization with additions of OM or PyOM (Figures 1 and 2). Simultaneously, increases in nSOC mineralization were greater with additions of OM than PyOM. These results are consistent with the idea that the activity of such microbial communities are more likely to be limited by C availability, such that the addition of PyOM could alleviate this constraint, resulting in general increased microbial activity, and, thus, increased SOC mineralization. In particular, the already low-C Quartzipsamment from Florida was especially vulnerable to increased nSOC losses with amendments. Although the Haplocalcid and Fragiudept soils also tended toward increased nSOC losses with the addition of PyOM, the Quartzipsamment was the only soil for which this effect was statistically significant for PyOM additions. These findings are consistent with previous studies across a range of soils and SOC contents (19–22). However, it is important to note that numerous other mechanisms could also contribute meaningfully to increased nSOC mineralization with organic amendments, as observed in other systems (17, 19, 20) and described in the introduction. However, we do not believe the effects we observed were primarily driven by pH shifts: the pH of four of the five soils were very similar (5.0-5.2). Additionally, we do not believe the effects were driven primarily by effects of the amendments on moisture: we adjusted moisture individually for each treatment. We do not believe that the effects were driven primarily by alleviation of a nutrient constraint with the addition of PyOM: the PyOM had relatively low N, and, furthermore, previous studies have often shown that soil CO_2_ emissions are inhibited by mineral N additions (54). Additionally, although the strongly-responding Quartzipsamment had the lowest measured mineral nutrients (Ca, Mg, and K; Table 1), the highest/second-highest nutrient soil was the Fragiudept, and it had the next strongest CO_2_ response to PyOM and OM amendments, suggesting that nutrient alleviation with PyOM or OM additions was not the dominant mechanism driving our observed effects.

On the one hand, the fact that the amendments had the least effect on the high-C soils suggests that, overall, the effects of increased nSOC mineralization with PyOM amendments might be less concerning, since the highest-C soils are less responsive. On the other hand, one might interpret it as being more concerning, since soils with the lowest SOC and lowest microbial activity to begin with, are most at risk for increased nSOC losses with PyOM amendments. This raises the question of which soils would be the best candidates for OM or PyOM additions. High-C soils seem to be lower risks for short-term increased CO_2_ emissions. However, other benefits to low-C soils, such as changes to water holding capacity, or total SOC content (PyOM-C + SOC), might outweigh this trade-off.

Even though our results strongly support the finding that short-term increases in CO_2_ emissions are most likely in soils with low C and/or low mineralization rates to begin with, it is important to note that these effects were observed over the *short term* – *i.e.*, over just a few weeks. As in other studies, the time period during which amendments increased net nSOC-derived CO_2_ emissions, the net increase usually began to level off, or even begin to decrease. Given this observation, and since other studies have observed net negative effects of PyOM amendments over longer time periods (16, 17), the findings from this study should be considered primarily within the context of short-term response to amendments.

### Magnitude of microbial community composition change mirrors magnitude of increases in nSOC-CO_2_

In response to our second question, our findings were also consistent with our primary hypothesis: we found that the degree to which soil microbial communities change with PyOM or OM amendments reflected the degree to which nSOC mineralization also increased (Figure 4). This supports the idea that the taxa that respond positively to PyOM and especially OM additions may also be the same taxa that are responsible for increased nSOC mineralization with PyOM or OM additions. Thus, a stronger shift toward these groups is accompanied by a stronger effect on nSOC mineralization. That said, it is important to note that, because we did not directly trace the fate of the organic substances into taxon-specific microbial biomass (*e.g.*, using an approach such as stable isotope probing), we have not conclusively demonstrated that the microbes that increased in abundance with additions were also the ones that metabolized the greater amount of nSOC. Still, it is not unreasonable to expect that increased total abundances of specific bacterial taxa might be accompanied by their increased activity as well. Overall, OM additions resulted in both a larger change in community composition, and also a larger increase in nSOC mineralization than did PyOM additions.

### PyOM responders differ across soils and do not reflect a common “charosphere”

Although PyOM additions did have a significant effect on microbial community composition, PyOM-induced changes in community composition were much smaller than the differences in community composition between different soils (Figure 3; Supplemental Tables S5 and S7; Supplemental Figures S1 and S2). Thus, PyOM did not result in a community dominated by the “charosphere” (33, 34), but, rather, resulted in detectable but relatively subtle shifts within a few of the existing taxa (Figures 5 and 6). We made a similar observation in our recent cross-study comparison of the effects of PyOM additions on soil bacterial community composition (27). This current study substantially improves our confidence in that observation, since it is not constrained by the challenges of cross-study differences in methods and materials and spans five different soils. Together, these observations underscore the importance of considering the effects of PyOM within the unique context of a given soil, rather than generalizing the effects of PyOM on soil microbial communities across all soils.

We were also interested in the specific taxa that responded to PyOM additions. In a previous field trial with the same Fragiudept soil and similar amendments (25), we identified a number of “common responders” to PyOM and OM after 82 days in the field. We suggested that those taxa may be most likely responsible for the short-term C mineralization effects of PyOM additions, and predicted that we would observe a similar phenomenon in the current study, possibly even across soils. This general trend persisted (Figure 6), in that OTUs that responded (positively or negatively) to one amendment tended to respond similarly to the other. Although there are a few taxa that are exceptions to this (respond positively to one amendment but negatively to the other), we hesitate to dwell too much on this response, since they tend to be low-abundance taxa to begin with. Because the same taxa that respond to PyOM over the short term also responded positively to OM, we suggest that this supports the idea that PyOM-responsive taxa in this study were likely responding to the small fraction of easily-mineralizable PyOM-C, and supporting the idea that a responsive fraction of the overall community might be responsible for short term increases in nSOC mineralization with PyOM amendments. Over longer timescales, we might expect different results as other mechanisms emerge. However, we were not necessarily able to identify a “core set” or PyOM responders across different soils. This is likely due in part to the small response overall to PyOM in the higher-C soils, and also to the diversity of organisms between soils. While there were 162 different PyOM-responsive OTUs, the same OTUs were often not present in the different soils: 62% of all 16S OTUs were detected (regardless of abundance) in only a single unamended soil (97% for ITS2), and 26% of all 16S OTUS were detected in only two different soils (2% for ITS2). In particular, since we used the dada2 OTU-picking algorithm, which can differentiate OTUs that differ by a single base pair, or “amplicon sequence variants”, it may be useful to consider common responders at a coarser phylogenetic scale. If we consider the OTUs at the genus level, there were numerous bacterial genera with OTUs that were responsive to PyOM in multiple soils, as well as OM amendments, as described in the results section. Some of the genera with PyOM-responsive OTUs across more than one soil were also identified as having PyOM-responsive OTUs in multiple studies in our previous meta-analysis, including *Flavisolibacter*, *Microvirga*, and *Noviherbaspirilllum* (27). Additionally, some of these PyOM-responsive bacteria are from genera that have been identified as being fire-responsive in other studies (*e.g.*, *Microvirga* (55), *Bacillus* (56), and *Noviherbaspirillum* (57)). Because all of the named taxa were also responsive to OM amendments over the short term, we raise the question of whether these OTUs may be responding to the more easily-mineralizable fractions of PyOM, or, in the case of fires, also to fire-released OM. Together, these taxa could represent interesting candidates for future investigation of the ecology of fire- and PyOM-responsive bacteria.

## Conclusions

While our short-term incubation indicates that low-C soils might be at the greatest risk for short-term C losses with OM or PyOM amendments, we note that the losses were greater with OM than with PyOM additions, and that many studies have shown that these short-term effects are relatively limited, and often even become net C increases over longer timescales. Together, our findings indicate that changes in microbial community composition mirrored changes in nSOC mineralization. This suggests that it may be likely that the change in CO_2_ emissions with the addition of amendments is governed by a specific subset of the microbial community, rather than a general stimulation of the entire community. Although these specific responsive organisms were not consistent across all soils, and depend on the native microbial community, certain taxa were identified as common responders. Future research could utilize techniques such as stable isotope probing to conclusively demonstrate which microbes are using the amendments as a C source, and to expand the research to more soil types, different timescales, and different PyOM materials to begin to develop a more comprehensive understanding of the specific microbial responders. It would also be interesting to determine whether or when our observation does not hold – whether there are conditions under which large community changes in response to organic amendments are not accompanied by changes in nSOC-CO_2_ emissions, and, conversely, whether there are conditions where large changes in CO_2_ emissions are observed, but not accompanied by changes in microbial community composition.

## Author Contribution Statement

T.W. and J.L. were responsible for the experimental design. T.W., S.D., K.H., A.E., and J.L. developed and optimized the experimental conditions. S.D., K.H., and A.E. set up and ran the in-lab experiment. T.W. and J.W. performed the DNA extractions, sequencing, and microbial bioinformatics. T.W. analyzed the data and T.W. and J.L. interpreted the data. T.W. drafted the manuscript and all authors contributed to, read, and approved the manuscript.

## Acknowledgements

We thank Dominic Woolf, who was instrumental in helping develop the original R script that was modified to analyze these data. The National Ecological Observatory Network is a program sponsored by the National Science Foundation and operated under cooperative agreement by Battelle Memorial Institute. This material is based in part upon work supported by the National Science Foundation through the NEON Program. We thank the NEON project, including Dave Tazik, Lee Stanish, and Samantha Weintraub for helping us access the NEON soils. We also would like to acknowledge the following individuals and organizations for providing soils for an initial planned version of the experiment which were, unfortunately, ultimately lost when a freezer broke down: John Anderson and Bernice Gamboa and the Jornada Experimental Range; William J. McShea and the Smithsonian Conservation Biology Institute; Mitch McClaran and the Santa Rita Experimental Range; Petra Royston, Ben Gottloeb, and Robert Nelson and the Disney Wilderness Preserve; Audrey Barker-Plotkin and the Harvard Forest; Stephen Coates and the Ordway Swisher Biological Station; Isabel Ashton, Tracey Baldwin and Craig Emerson and the Rocky Mountain National Park, Central Plains Experimental Range, and the North Sterling NEON sites; Holly Johnson, Matt Sanderson, and Mark Liebig and the Northern Great Plains Research Lab; Ty Lindberg at NEON; all others who helped collect and ship those soils. An NSF Doctoral Dissertation Improvement Grant award to T.W. and J.L. supported the bulk of this research [NSF-1406195]; U.S. Department of Energy also helped support T.W. [DE-SC0016365].

## Supplementary Information

Supplementary information is available at X.

## References

1. Ciais P, Sabine C, Bala G, Bopp L, Brovkin V, Canadell J, Chhabra A, DeFries R, Galloway J, Heimann M, Jones C, Le Quéré C, Myneni RB, Piao S, Thornton P. 2013. Carbon and other biogeochemical cycles, pp. 465–570. *In* Stocker, TF, Quin, D, Plattner, GK, Tignor, M, Allen, SK, Boschung, J, Nauels, A, Xia, Y, Bex, V, Midgley, PM, P (eds.), Climate Change - The Physical Science Basis. Contribution of Working Group I to the Fifth Assessment Report of the Intergovernmental Panel on Climate Change. Cambridge Univ Press, Cambridge, UK.

2. Schmidt MWI, Torn MS, Abiven S, Dittmar T, Guggenberger G, Janssens IA, Kleber M, Kögel-Knabner I, Lehmann J, Manning DAC, Nannipieri P, Rasse DP, Weiner S, Trumbore SE. 2011. Persistence of soil organic matter as an ecosystem property. Nature 478:49–56.

3. Zimmerman AR, Mitra S. 2017. Trial by Fire: On the Terminology and Methods Used in Pyrogenic Organic Carbon Research. Frontiers in Earth Science 5:354.

4. Jones MW, Santín C, Van Der Werf GR, Doerr SH. 2019. Global fire emissions buffered by the production of pyrogenic carbon. Nature Geoscience, 12:742–747.

5. Reisser M, Purves RS, Schmidt MWI, Abiven S. 2016. Pyrogenic Carbon in Soils: A Literature-Based Inventory and a Global Estimation of Its Content in Soil Organic Carbon and Stocks. Frontiers in Earth Science, 4:80.

6. Lehmann J. 2007. A handful of carbon. Nature 447:143–144.

7. Laird DA. 2008. The charcoal vision: A win–win–win scenario for simultaneously producing bioenergy, permanently sequestering carbon, while improving soil and water quality. Agronomy Journal 100:178–181.

8. Wardle DA, Nilsson MC, Zackrisson O. 2008. Fire-Derived Charcoal Causes Loss of Forest Humus. Science 320:629–629.

9. Wardle DA, Nilsson MC, Zackrisson O. 2008. Response to Comment on “Fire-Derived Charcoal Causes Loss of Forest Humus.” Science 321:1295d–1295d.

10. Lehmann J, Sohi S. 2008. Comment on “Fire-Derived Charcoal Causes Loss of Forest Humus.” Science 321:1295c–1295c.

11. Zimmerman AR, Bin Gao, Ahn M-Y. 2011. Positive and negative carbon mineralization priming effects among a variety of biochar-amended soils. Soil Biology and Biochemistry 43:1169–1179.

12. Kuzyakov Y, Subbotina I, Chen H, Bogomolova I, Xu X. 2009. Black carbon decomposition and incorporation into soil microbial biomass estimated by ^14^C labeling. Soil Biology and Biochemistry 41:210–219.

13. Luo Y, Durenkamp M, De Nobili M, Lin Q, Brookes PC. 2011. Short term soil priming effects and the mineralisation of biochar following its incorporation to soils of different pH. Soil Biology and Biochemistry 43:2304–2314.

14. Bingeman CW, Varner JE, Martin WP. 1953. The effect of the addition of organic materials on the decomposition of an organic soil. Soil Science Society of America Journal 17:34–38.

15. Blagodatskaya E, Kuzyakov Y. 2008. Mechanisms of real and apparent priming effects and their dependence on soil microbial biomass and community structure: critical review. Biology and Fertility of Soils 45:115–131.

16. Whitman T, Zhu Z, Lehmann J. 2014. Carbon Mineralizability Determines Interactive Effects on Mineralization of Pyrogenic Organic Matter and Soil Organic Carbon. Environmental Science & Technology 48:13727–13734.

17. DeCiucies S, Whitman T, Woolf D, Enders A, Lehmann J. 2018. Priming mechanisms with additions of pyrogenic organic matter to soil. Geochimica et Cosmochimica Acta 238:329–342.

18. Cheng H, Hill PW, Bastami MS, Jones DL. 2017. Biochar stimulates the decomposition of simple organic matter and suppresses the decomposition of complex organic matter in a sandy loam soil. GCB Bioenergy 9:1110–1121.

19. Maestrini B, Nannipieri P, Abiven S. 2014. A meta-analysis on pyrogenic organic matter induced priming effect. GCB Bioenergy 7:577–590.

20. Whitman T, Singh BP, Zimmerman AR. 2015. Priming effects in biochar-amended soils: Implications of biochar-soil organic matter interactions for carbon storage, pp. 455–487. *In* Lehmann, J, Joseph, S (eds.), Biochar for Environmental Management, 2nd ed. Routledge, New York, NY.

21. Wang J, Xiong Z, Kuzyakov Y. 2016. Biochar stability in soil: meta-analysis of decomposition and priming effects. GCB Bioenergy 8:512–523.

22. Ding F, van Zwieten L, Zhang W, Weng ZH, Shi S, Wang J, Meng J. 2018. A meta-analysis and critical evaluation of influencing factors on soil carbon priming following biochar amendment. Journal of Soils and Sediments 18:1507–1517.

23. Woolf D, Lehmann J. 2012. Modelling the long-term response to positive and negative priming of soil organic carbon by black carbon. Biogeochemistry 111:83–95.

24. Woolf D, Amonette JE, Street-Perrott FA, Lehmann J, Joseph S. 2010. Sustainable biochar to mitigate global climate change. Nature Communications 1:1–9.

25. Whitman T, Pepe-Ranney C, Enders A, Koechli C, Campbell A, Buckley DH, Lehmann J. 2016. Dynamics of microbial community composition and soil organic carbon mineralization in soil following addition of pyrogenic and fresh organic matter. The ISME Journal 10:2918–2930.

26. Yu Z, Chen L, Pan S, Li Y, Kuzyakov Y, Xu J, Brookes PC, Luo Y. 2018. Feedstock determines biochar-induced soil priming effects by stimulating the activity of specific microorganisms. European Journal of Soil Science 69:521–534.

27. Woolet J, Whitman T. Review: Pyrogenic organic matter effects on soil bacterial community composition. Soil Biology and Biochemistry 141:107678.

28. Dai Z, Hu J, Xu X, Zhang L, Brookes PC, He Y, Xu J. 2016. Sensitive responders among bacterial and fungal microbiome to pyrogenic organic matter (biochar) addition differed greatly between rhizosphere and bulk soils. Scientific Reports 1–11.

29. Nielsen S, Minchin T, Kimber S, van Zwieten L, Gilbert J, Munroe P, Joseph S, Thomas T. 2014. Comparative analysis of the microbial communities in agricultural soil amended with enhanced biochars or traditional fertilisers. Agriculture, Ecosystems and Environment 191:73–82.

30. Wu H, Zeng G, Liang J, Chen J, Xu J, Dai J, Li X, Chen M, Xu P, Zhou Y, Li F, Hu L, Wan J. 2016. Responses of bacterial community and functional marker genes of nitrogen cycling to biochar, compost and combined amendments in soil. Applied Microbiology and Biotechnology 100:8583–8591.

31. Imparato V, Hansen V, Santos SS, Nielsen TK, Giagnoni L, Hauggaard-Nielsen H, Johansen A, Renella G, Winding A. 2016. Gasification biochar has limited effects on functional and structural diversity of soil microbial communities in a temperate agroecosystem. Soil Biology and Biochemistry 99:128–136.

32. Yao Q, Liu J, Yu Z, Li Y, Jin J, Liu X, Wang G. 2017. Changes of bacterial community compositions after three years of biochar application in a black soil of northeast China. Applied Soil Ecology 113:11–21.

33. Quilliam RS, Glanville HC, Wade SC. 2013. Life in the “charosphere”–Does biochar in agricultural soil provide a significant habitat for microorganisms? Soil Biology and Biochemistry 65:287–293.

34. Clough TJ, Condron LM. 2010. Biochar and the Nitrogen Cycle: Introduction. Journal of Environment Quality 39:1218.

35. National Ecological Observatory Network. 2016. Soil microbe prototype 16S sequence data, 2009–2010.

36. Whitman T, Lehmann J. 2015. A dual-isotope approach to allow conclusive partitioning between three sources. Nature Communications 6:8708.

37. Walters W, Hyde ER, Berg-Lyons D, Ackermann G, Humphrey G, Parada A, Gilbert JA, Jansson JK, Caporaso JG, Fuhrman JA, Apprill A, Knight R. 2015. Improved Bacterial 16S rRNA Gene (V4 and V4-5) and Fungal Internal Transcribed Spacer Marker Gene Primers for Microbial Community Surveys. mSystems 1:e00009–15.

38. Taylor DL, Walters WA, Lennon NJ, Bochicchio J, Krohn A, Caporaso JG, Pennanen T. 2016. Accurate Estimation of Fungal Diversity and Abundance Through Improved Lineage-Specific Primers Optimized for Illumina Amplicon Sequencing. Applied and Environmental Microbiology 82:AEM.02576–16–7226.

39. Kozich JJ, Westcott SL, Baxter NT, Highlander SK, Schloss PD. 2013. Development of a Dual-Index Sequencing Strategy and Curation Pipeline for Analyzing Amplicon Sequence Data on the MiSeq Illumina Sequencing Platform. Applied and Environmental Microbiology 79:5112–5120.

40. Callahan BJ, McMurdie PJ, Rosen MJ, Han AW, Johnson AJA, Holmes SP. 2016. DADA2: High-resolution sample inference from Illumina amplicon data. Nature Methods 13:581–583.

41. Wang Q, Garrity GM, Tiedje JM, Cole JR. 2007. Naive Bayesian classifier for rapid assignment of rRNA sequences into the new bacterial taxonomy. Applied and Environmental Microbiology 73:5261–5267.

42. Quast C, Pruesse E, Yilmaz P, Gerken J, Schweer T, Yarza P, Peplies J, Glöckner FO. 2013. The SILVA ribosomal RNA gene database project: improved data processing and web-based tools. Nucleic Acids Research 41:D590–D596.

43. Bolyen E, Rideout JR, Dillon MR, Bokulich NA, Abnet CC, Al-Ghalith GA, Alexander H, Alm EJ, Arumugam M, Asnicar F, Bai Y, Bisanz JE, Bittinger K, Brejnrod A, Brislawn CJ, Brown CT, Callahan BJ, Caraballo-Rodríguez AM, Chase J, Cope EK, Da Silva R, Diener C, Dorrestein PC, Douglas GM, Durall DM, Duvallet C, Edwardson CF, Ernst M, Estaki M, Fouquier J, Gauglitz JM, Gibbons SM, Gibson DL, Gonzalez A, Gorlick K, Guo J, Hillmann B, Holmes S, Holste H, Huttenhower C, Huttley GA, Janssen S, Jarmusch AK, Jiang L, Kaehler BD, Bin Kang K, Keefe CR, Keim P, Kelley ST, Knights D, Koester I, Kosciolek T, Kreps J, Langille MGI, Lee J, Ley R, Liu Y-X, Loftfield E, Lozupone C, Maher M, Marotz C, Martin BD, McDonald D, McIver LJ, Melnik AV, Metcalf JL, Morgan SC, Morton JT, Naimey AT, Navas-Molina JA, Nothias LF, Orchanian SB, Pearson T, Peoples SL, Petras D, Preuss ML, Pruesse E, Rasmussen LB, Rivers A, Robeson MS, Rosenthal P, Segata N, Shaffer M, Shiffer A, Sinha R, Song SJ, Spear JR, Swafford AD, Thompson LR, Torres PJ, Trinh P, Tripathi A, Turnbaugh PJ, Ul-Hasan S, van der Hooft JJJ, Vargas F, Vázquez-Baeza Y, Vogtmann E, Hippel von M, Walters W, Wan Y, Wang M, Warren J, Weber KC, Williamson CHD, Willis AD, Xu ZZ, Zaneveld JR, Zhang Y, Zhu Q, Knight R, Caporaso JG. 2019. Reproducible, interactive, scalable and extensible microbiome data science using QIIME 2. Nature Biotechnology 37:852–857.

44. Zhang J, Kobert K, Flouri T, Stamatakis A. 2014. PEAR: a fast and accurate Illumina Paired-End reAd mergeR. Bioinformatics 30:614–620.

45. R Core Team. R: A language and environment for statistical computing. ISBN 3-900051-07-0. Vienna, Austria.

46. Balesdent J, Mariotti A. 1996. Measurement of soil organic matter turnover using ^13^C natural abundance., pp. 83–111. *In* Boutton, TW, Yamasaki, SI (eds.), Mass Spectrometry of Soils.

47. McMurdie PJ, Holmes S. 2013. phyloseq: An R Package for Reproducible Interactive Analysis and Graphics of Microbiome Census Data. PLoS ONE 8:e61217.

48. Wickham H. ggplot2: Elegant Graphics for Data Analysis. Springer-Verlag New York.

49. Wickham H, François R, Henry L, Müller K. dplyr: A Grammar of Data Manipulation.

50. Bray JR, Curtis JT. 1957. An Ordination of the Upland Forest Communities of Southern Wisconsin. Ecological Monographs 27:325–349.

51. Legendre P, Gallagher ED. 2001. Ecologically meaningful transformations for ordination of species data. Oecologia 129:271–280.

52. Oksanen J, Blanchet FG, Kindt R, Legendre P, Minchin PR, OHara RB, Simpson GL, Solymos P, Stevens MHH, Wagner H. vegan: Community Ecology Package, 2nd ed.

53. Martin BD. corncob: Count Regression for Correlated Observations with the Beta-Binomial.

54. Ramirez KS, Craine JM, Fierer N. 2012. Consistent effects of nitrogen amendments on soil microbial communities and processes across biomes. Global Change Biology 18:1918–1927.

55. Fernández-González AJ, Martínez-Hidalgo P, Cobo-Díaz JF, Villadas PJ, Martínez-Molina E, Toro N, Tringe SG, Fernández-López M. 2017. The rhizosphere microbiome of burned holm-oak: potential role of the genus *Arthrobacter* in the recovery of burned soils. Scientific Reports 7:6008.

56. Cobo-Díaz JF, Fernández-González AJ, Villadas PJ, Robles AB, Toro N, Fernández-López M. 2015. Metagenomic Assessment of the Potential Microbial Nitrogen Pathways in the Rhizosphere of a Mediterranean Forest After a Wildfire. Microbial Ecology 69:895–904.

57. Whitman T, Whitman E, Woolet J, Flannigan MD, Thompson DK, Parisien M-A. 2019. Soil Bacterial and Fungal Response to Wildfires in the Canadian Boreal Forest Across a Burn Severity Gradient. Soil Biology and Biochemistry, 138:107571.

